# Loss of phosphorylation sites required for β-arrestin recruitment in the calcium-sensing receptor cytoplasmic region may contribute to hypocalcemia

**DOI:** 10.64898/2026.04.21.719673

**Authors:** Rachael A. Wyatt, Hassan Massalha, Caroline M. Gorvin

## Abstract

The calcium-sensing receptor (CaSR) is a class C G protein-coupled receptor (GPCR) with an important role in calcium homeostasis, activating mutations of which cause hypocalcemia. Despite the receptors clinical importance and evidence suggesting β-arrestin-1/2 can modulate CaSR signaling, the residues within CaSR required for β-arrestin recruitment and the role of β-arrestin in desensitization, trafficking and signaling are ill defined. We confirmed that β-arrestin-1 and β-arrestin-2 are recruited to CaSR upon receptor activation. Deletion of the distal cytoplasmic region of CaSR, which replicates several mutations identified in individuals with hypocalcemia, reduced β-arrestin-1/2 recruitment and enhanced receptor signaling. Examination of the receptor cytoplasmic region identified three regions of serine and threonine residues that resemble phosphorylation codes identified in other GPCRs. Mutation of each of these residues to alanine demonstrated one region between amino acids 1003-1011 is important for both β-arrestin-1 and β-arrestin-2 recruitment, while disruption of a second region between residues 976-981 impaired β-arrestin-2 recruitment. Alanine mutagenesis of these residues also reduced CaSR signaling and impaired receptor internalization suggesting an important role in receptor desensitization. Three variants in the ClinVar dataset reported in disorders of calcium homeostasis were identified and two, T1006M and T1008P, were shown to reduce β-arrestin recruitment and enhance CaSR signaling. Further analysis of the T1006M variant showed it reduced CaSR internalization. Thus, we have identified two regions within the CaSR cytoplasmic region that are important for β-arrestin recruitment, receptor desensitization and internalization. Mutation of residues within these regions may represent another mechanism for the pathogenesis of hypocalcemia.

## Introduction

The class C GPCR, CaSR, is essential for maintaining calcium homeostasis by regulating parathyroid hormone release and urinary calcium excretion (1). Receptor activation elicits signaling predominantly by G_q/11_, stimulating phospholipase C to increase IP_3_-mediated intracellular calcium release (Ca^2+^_i_) and mitogen-activated protein kinase (MAPK) pathways (e.g. extracellular signal-regulated kinases (ERK1/2)), and G_i/o_ to reduce cAMP (1,2). For most GPCRs, activation is followed by recruitment of GPCR kinases (GRKs) that phosphorylate the receptor at serine and threonine residues in the intracellular regions (intracellular loops (ICL) or C terminal domain (CTD)) to promote interactions with cytosolic β-arrestins (β-arrestin-1 and β-arrestin-2)(3–7). In the classical model of GPCR signaling, recruitment of β-arrestins physically disrupts G protein coupling and facilitates clathrin-mediated endocytosis of the receptor in a mechanism known as desensitization (3). However, multiple studies have shown that β-arrestins can also act as scaffolds for MAPK signaling (8–10), and persistent binding of β-arrestin to internalized GPCRs can produce endosomal activation of G protein to sustain receptor responses (11). Although CaSR can produce sustained G_q/11_ signals from endosomes (2,12), this is by a β-arrestin-independent pathway (12), which has also been demonstrated for other GPCRs (13). Moreover, it is unclear how important β-arrestin recruitment is for CaSR desensitization, signaling and internalization. There is some evidence that β-arrestins are recruited to CaSR and may affect signaling. CaSR coimmunoprecipitates with β-arrestin-1, which was dependent on interaction with the cytoplasmic region of CaSR (14). One study showed co-transfection of β-arrestins with GRK2 and GRK4 reduced CaSR-mediated activity (14), while another showed PKC phosphorylation of sites within the cytoplasmic tail are crucial for β-arrestin recruitment and termination of IP3 signaling (15). β-arrestin-1 may also have distinct scaffolding roles in CaSR-mediated cytoskeletal remodeling (16) However, CaSR-mediated phosphorylation of ERK1/2 has been shown to be independent of β-arrestin-2 in one study (17), while another describes its importance for ERK1/2 signaling (18).

Similarly, the importance of β-arrestin recruitment for CaSR internalization has been subject to debate. Studies in HEK293 overexpressing CaSR in the early 2000s showed agonist-mediated internalization of CaSR was not enhanced by co-transfection of β-arrestin-1, β-arrestin-2, GRK2 or GRK3, nor was there a change in cell surface expression (14,15), suggesting CaSR trafficking may not require β-arrestin coupling. Since these studies, there have been substantial developments in assays to assess receptor internalization and a 2019 study that utilized HEK293 depleted of β-arrestin-1 and β-arrestin-2 and a kinetic assay to measure internalization of SNAP-tagged CaSR demonstrated both constitutive and agonist-mediated endocytosis are dependent on β-arrestin-1 and β-arrestin-2 (19). However, they did not investigate the CaSR residues involved in β-arrestin recruitment.

There is significant evidence that different patterns of receptor phosphorylation (the phosphorylation barcode) can induce distinct β-arrestin structural conformations that govern how the receptor signals, is desensitized and internalized (3–7,20). Three phosphorylation motifs of different lengths have been described (PxxPxxP/E/D or PxPxxP/E/D or P-x-P-P, where P is a phosphorylated serine or threonine) and proposed to form an extensive network of electrostatic interactions with positively charged pockets in arrestin (6,20). NMR subsequently demonstrated that GPCR C-termini, which behave as intrinsically disordered regions with transient secondary structures, undergo structural transitions upon GRK phosphorylation to form β-strand conformations in localized segments corresponding to arrestin-interacting regions (7). Typically, class A GPCRs harbor one complete or partial phosphorylation code, while class B GPCRs have at least two phosphorylation codes, consistent with transient interactions between β-arrestins and class A GPCRs, and longer-lived interactions that promote distinct signaling patterns with class B receptors (6,21). Differential phosphorylation of some GPCRs has also been shown to direct signalling towards distinct pathways (i.e. they bias signaling)(22). The phosphorylation patterns of class C GPCRs are generally understudied although there is evidence that some members of the metabotropic glutamate receptor (mGluR) family have phosphorylation codes within their CTD (23).

The CaSR has a long CTD of over 200 amino acids (residues 865-1078). Biochemical studies have shown that truncations of the latter part of the CTD (after residue 886) can enhance receptor surface expression and downstream signalling (24–27). Consistent with this, several *CASR* truncation mutations that affect the distal CTD increase receptor activity and cause autosomal dominant hypocalcemia (ADH)(28,29), whereas inactivating truncation mutations that cause the opposite disorder, familial hypocalciuric hypercalcemia (FHH), cluster in the proximal CTD (30,31). Truncation of the distal CTD likely enhances signaling by a combination of mechanisms including retention of essential residues for signaling in the proximal CTD (32,33) as well as an ER checkpoint motif important for cell surface expression (26), whereas the distal CTD region harbours a dileucine motif required for receptor internalization and motifs that target CaSR to degradation pathways, which when lost will increase signaling (24,34). However, it is also possible that this distal CTD region also harbors phosphorylation sites that promote β-arrestin recruitment.

Here we investigated which residues in the CaSR intracellular region are important for β-arrestin recruitment. We identified two motifs in the distal CTD that may have roles in β-arrestin recruitment and showed clinically relevant variants within these sites can enhance signaling by reducing β-arrestin recruitment.

## Experimental Procedures

### Cell culture, transfection and plasmid constructs

Adherent HEK293 (AdHEK) cells were purchased from Agilent Technologies (Santa Clara, CA, USA). HEK293 cells with deletion of β-arrestin-1 and β-arrestin-2 (HEK-βarr1/2 KO) were described previously (35). AdHEK cells and β-arrestin-1/2 knockout cells were maintained in DMEM-Glutamax media (Sigma, Gillingham, UK) with 10% fetal bovine serum (FBS, Sigma) at 37°C, 5% CO_2_. Cells were maintained in penicillin-streptomycin-free media and routinely screened to ensure they were mycoplasma-free using the TransDetect Luciferase Mycoplasma Detection kit (Generon). Transfections were performed using Lipofectamine 2000 (Invitrogen, Waltham, MA).

The CaSR-SmC plasmid has been described (2), and the CaSR-SmC-895X truncation was generated by the same method. The NanoBiT expression plasmids, LgC-βarr1, LgC-βarr2 have been described (36) and were kindly provided by Asuka Inoue (Tohoku University, Sendai, Japan). The HA-SNAP-CaSR plasmid was generated within the pRK5-mGluR2 vector(37) (a gift from Joshua Levitz, Weill Cornell Medicine) and has an N-terminal signal peptide followed by the HA affinity tag, SNAP self-labeling protein tag capable of conjugation to organic dyes, and human CaSR (amino acids 20-1078). SmC-CaSR and HA-SNAP-CaSR mutants were generated using oligonucleotides from Sigma, the QuikChange Lightning site-directed mutagenesis kit (Agilent Technologies, Santa Clara, CA) and sequenced by Source Bioscience (Nottingham, UK).

### Bioinformatics Resources

Phosphorylation site predictions were made using the Phospho-ELM(38) and NetPhos3.1(39) resources. CADD (combined annotation dependent depletion)(40), SIFT (sorting intolerant from intolerant)(41) and REVEL (Rare Exome Variant Ensemble Learner)(42) were used to predict the effect of amino acid substitutions. Amino acid conservation was examined in CaSR mammalian orthologs using ClustalOmega(43). The ClinVar database(44) and the Human Gene Mutation Database (https://www.hgmd.cf.ac.uk/ac/index.php) were examined to identify missense variants in the CaSR cytoplasmic region.

### Detection of ARRB1 and ARRB2 in published parathyroid scRNA-seq datasets

Single-cell RNA-sequencing datasets containing parathyroid cells from healthy (non-parathyroid adenoma) human adult (45,46), and Macaca mulatta (46) samples were included in this analysis. We followed default scanpy integration steps including all cells per study from human adult and Macaca mulatta. Briefly, basic filtering was applied to post-QC raw count data, retaining cells with more than 500 detected genes and genes expressed in more than 3 cells. Highly variable genes were then identified using scanpy (v1.11.5) ‘highly_variable_genes’ (flavor = ‘seurat_v3’, n_top_genes = 2000). The data for each study were integrated by sample using Harmony (47) (harmonypy; v0.2.0, theta = 0) with the number of PCA dimensions selected based on an elbow plot. For downstream analysis, we subset parathyrocyte clusters based on presence of the canonical marker genes *PTH, GCM2, GATA3* and *CASR* and assigned a cell-cycle phase to each cell using a cycling gene list from previous studies (48) with the scanpy ‘score_genes_cell_cycle’ function. Normalized expression values are shown per study for non-cycling (G1) cells and cycling (S and G2M) cells.

### Assessment of cell surface expression

For assessment of surface expression of HA-SNAP-CaSR plasmids, AdHEK cells were seeded at 10,000 cells/well in black 96-well plates and transfected the same day with 100 ng pcDNA or 100 ng HA-SNAP-CaSR (WT or variants). Cells were labelled 48-hours later with SNAP-647, then fluorescence read on a Glomax plate reader (Promega) following three washes with PBS. Expression levels were normalized to pcDNA control transfected cells. For assessment of CaSR-895X cell surface expression, AdHEK cells were transfected with 100ng CaSR-SmC WT or 895X plasmid and cells fixed 48-hours later in 4% PFA (Fisher Scientific UK) in PBS, then labelled with the Mouse monoclonal anti-CaSR primary antibody (1:1000, clone ADD/5C10, Abcam, Cat#ab19347, RRID: AB_444867), followed by Alexa Fluor 647 donkey anti-mouse secondary antibody (Abcam Catalog. number ab181292, RRID:AB_3351687). Cells were washed, then fluorescence read on a Glomax plate reader.

### Western blot analysis

Cells were seeded into 6-well plates then transfected 24-hours later with HA-SNAP-CaSR wild-type or mutant protein at 1µg per well. Cells were lysed after 48-hours in NP40 buffer and western blot analysis performed, as described (2). Blots were blocked in 5% marvel/TBS-T, then probed with the primary antibody anti-CaSR (1:1000, clone ADD/5C10, Abcam, Cat#ab19347, RRID: AB_444867)and the Goat Anti-Mouse IgG HRP Conjugated secondary antibody (1:3000, Catalog. no. 1706516, BioRad, RRID:AB_2921252). After visualizing and stripping, the blot was reprobed with the rabbit anti-calnexin polyclonal antibody (1:1000, rabbit polyclonal, Catalog number AB2301, Millipore, RRID:AB_10948000), followed by the Goat Anti-Rabbit IgG, HRP Conjugated secondary antibody (1:3000, Sigma, Catalog no. A0545; RRID:AB_257896).

For pERK1/2 studies, cells were transfected with 1μg plasmid and 36 hours later media was replaced with Ca^2+^ and Mg^2+^-free HBSS. At least 30 minutes later, cells were stimulated with 5mM Ca^2+^ for 5 or 10 minutes, then cells lysed and western blot performed. After analysis for pERK1/2 (rabbit polyclonal, Catalog no. 9101L, Cell Signaling Technologies, RRID:AB_331646), blots were stripped and re-probed with an anti-total ERK1/2 antibody (rabbit polyclonal, Catalog no. 4695S, Cell Signaling Technologies, RRID:AB_390779), then either rabbit anti-calnexin or rabbit anti-alpha-tubulin monoclonal primary antibody (Abcam, Catalog no. ab176560, RRID:AB_2860019) as loading controls. Blots were visualized using the Clarity Western ECL kit (BioRad) on a BioRad Chemidoc XRS+ system. Densitometry was performed using ImageJ (NIH), and protein quantities normalized to calnexin for CaSR and to total ERK1/2 for pERK1/2 blots.

### NanoBiT β-arrestin recruitment assay

NanoBiT recruitment assays were performed using methods adapted from those previously described (36,49). NanoBiT β-arrestin assays were performed in HEK-βarr1/2 KO cells. Cells were plated in six-well plates and transfected 48 hours later with 500ng CaSR-SmC wild-type or mutant plasmids with 200ng LgC-β-arrestin-1 or LgC-β-arrestin-2.

Following 48 hours, cells were harvested in DMEM-FluoroBrite (Gibco, Waltham, MA) with 10% FBS and 4mM L-glutamine (Gibco) and seeded in 8 wells of a 96-well plate. Media was changed to calcium- and magnesium-free Hank’s Balanced Saline Solution (HBSS, Sigma) approximately 60 minutes before the assay was performed. NanoGlo reagent (Promega) was added 15 minutes prior to assay performance to stabilize baseline reads. Luminescence values were performed on a Promega Glomax at 37°C. Agonist (CaCl_2_, VWR) was prepared in HBSS at 10x concentration and added to wells once baselines had stabilized, and responses recorded immediately after agonist addition for 35 minutes. Response values at each concentration were calculated by subtracting the agonist luciferase values from a vehicle-treated sample for each CaSR plasmid, as previously described (50).

Data was plotted using GraphPad Prism 7. The area-under-the-curve (AUC) was calculated at each agonist concentration, then the maximal wild-type response set as 100%, and all mutant responses normalized to this value to compare wild-type and mutant responses in each experiment. Nonlinear regression of concentration-response curves was performed with GraphPad Prism using the normalized response at each [Ca^2+^]_e_ for the determination of the mean half-maximal concentration (EC_50_).

### Structured illuminated microscopy (SIM)

SIM was performed as previously described. Cells were plated on 24mm coverslips (VWR) and transfected with 500ng of each plasmid 36-hours prior to experiments. Cells were incubated in Ca^2+^-and Mg^2+^-free HBSS for at least one hour, then exposed to 1:1000 anti-HA mouse monoclonal antibody (BioLegend Cat#901514, RRID:AB_2565336) with either vehicle or 5 mM CaCl_2_ for 30 minutes. Cells were fixed in 4% PFA/PBS, permeabilized with 1% Triton-X100/PBS and exposed to the anti-mouse IgG (H+L), F(ab’)2 fragment (Alexa Fluor® 647 Conjugate) secondary antibody (1:1000, Cell Signaling Technology, Catalog. no. 4410; RRID:AB_1904023). Samples were imaged on a Nikon N-SIM system (Ti-2 stand, Cairn TwinCam with 2 × Hamamatsu Flash 4 sCMOS cameras, Nikon laser bed 488 and 647 nm excitation lasers, Nikon 100 × 1.49 NA TIRF Apo oil objective). SIM data was reconstructed using NIS-Elements (v. 5.21.03) slice reconstruction. Vesicles were counted in ImageJ using previously described protocols (51). A manual threshold was set for each image to subtract background. Images were processed using ‘Process>Binary>Convert to mask’ and ‘Analyze particles’ in ImageJ. Images were manually inspected to ensure vesicles were captured by this method and observers were blinded to the experimental conditions. The size of each cell was measured in ImageJ and the number of vesicles normalized to the cell area.

### Confocal microscopy

AdHEK cells were plated on coverslips in six-well plates and transiently transfected with 1µg of either the CaSR-SmC WT or 895X plasmid. Cells were fixed in 4% PFA/PBS, permeabilized with 1% Triton-X100/PBS and exposed to the primary mouse monoclonal anti-CaSR primary antibody (1:1000, clone ADD/5C10, Abcam, Cat#ab19347, RRID: AB_444867), then the secondary antibody Donkey anti-mouse IgG Alexa Fluor-488 (1:1000, Molecular Probes, Cat# A-21202, RRID: AB_141607). Cells were mounted in Prolong Gold Antifade reagent (Invitrogen) and images captured using a Zeiss LSM780 confocal microscope with a Plan-Apochromat x63/1.2/water DIC objective. An argon laser (488 nm) was used to excite Alexa Fluor-488.

### HILO microscopy

AdHEK cells were seeded on 24mm coverslips (VWR) and transfected with 500ng of each plasmid 24-hours prior to experiments. On the day of the experiment SNAP-Surface Alexa Fluor 647 (NEB) was diluted 1:1000 in FluoroBrite complete media and applied to cells for 20-minutes, before washing and imaging. HILO images were acquired on a custom-built TIRF microscope (Cairn Research) containing an Eclipse Ti2 (Nikon) equipped with an EMCCD camera (iXon Ultra, Andor), a 488 nm diode laser, a hardware Perfect Focus System, a TIRF iLas2 module, and a 100x oil-immersion objective (NA 1.49, Nikon). Coverslips were mounted onto plastic imaging chambers with a rubber seal and filled with imaging medium (Ca^2+^- and Mg^2+^-free HBSS with 10mM HEPES). The objective and samples were maintained at 37°C. Images were acquired on MetaMorph software (Molecular Devices) using a frame exposure of 50–200 ms with two images acquired before ligand stimulation (3mM Ca^2+^_e_) and a subsequent image taken every 30 seconds thereafter, up to 20 minutes. Images were analyzed using ImageJ.

### Fluo-4 intracellular calcium assays

AdHEK cells were plated in black sided 96-well plates and transfected with 50ng CaSR-SmC WT or mutant plasmids per well. Cells were incubated for 48 hours prior to performance of the Fluo-4 intracellular calcium assays. Fluo-4 AM (Molecular Probes, Life Technologies) was resuspended in DMSO/ 0.03% pluronic acid, then a working solution made on the day of the experiment at 1:1000 dilution in calcium- and magnesium-free HBSS (Sigma) with 10mM HEPES (Gibco) (HANKS-HEPES solution). Cells were incubated with 50µL Fluo-4 working solution and incubated at room temperature for one hour. Cells were washed once in HANKS-HEPES solution, then 50µL fresh HANKS-HEPES added prior to assay performance on a Glomax Discover at 37°C with excitation 494 nm and emission at 516 nm. Ten baseline reads were made per well prior to injection of drugs from preloaded injectors using an automated protocol. Following injection of each drug concentration, ten reads were performed. At least three technical replicates were performed per condition and the position of each condition varied across each plate. Data was plotted in GraphPad Prism. The maximal response for each drug concentration was derived, normalized to baseline responses of the CaSR-WT transfected wells and concentration-response curves plotted with a 4-parameter sigmoidal fit.

### Statistical Analysis

The number of experimental replicates denoted by *n* is indicated in each figure legend. Data were plotted and statistical analyses performed in Graphpad Prism 7. Normality tests (Shapiro–Wilk or D’Agostino–Pearson) were performed on all datasets to determine whether parametric or non-parametric tests were appropriate. A *P*-value of <0.05 was considered statistically significant. A minimum of three independently transfected replicates was performed in all cell-based assays, and each experiment was performed on separate occasions with different passages of cells.

## Results

### CaSR activation recruits β-arrestin-1 and β-arrestin-2 to the cell surface

We used the NanoBiT β-arrestin assay (36) to determine whether CaSR activation affects β-arrestin-1/2 recruitment in HEK293 cells depleted of β-arrestin-1/2 by CRISPR-Cas gene-editing. Cells were transfected with CaSR-SmC, which we have previously shown to express and signal similarly to FLAG-tagged CaSR(2), and LgC-β-arrestin-1 or LgC-β-arrestin-2 and exposed to Ca^2+^_e_ to stimulate the receptor. This showed robust luminescence responses for both β-arrestin-1 and β-arrestin-2 (Figure 1A). Responses were larger for β-arrestin-1, although the EC_50_ values were not significantly different (Figure 1B). Therefore, CaSR can recruit β-arrestin-1 and β-arrestin-2 when expressed in HEK293 cells.

**Figure 1.**
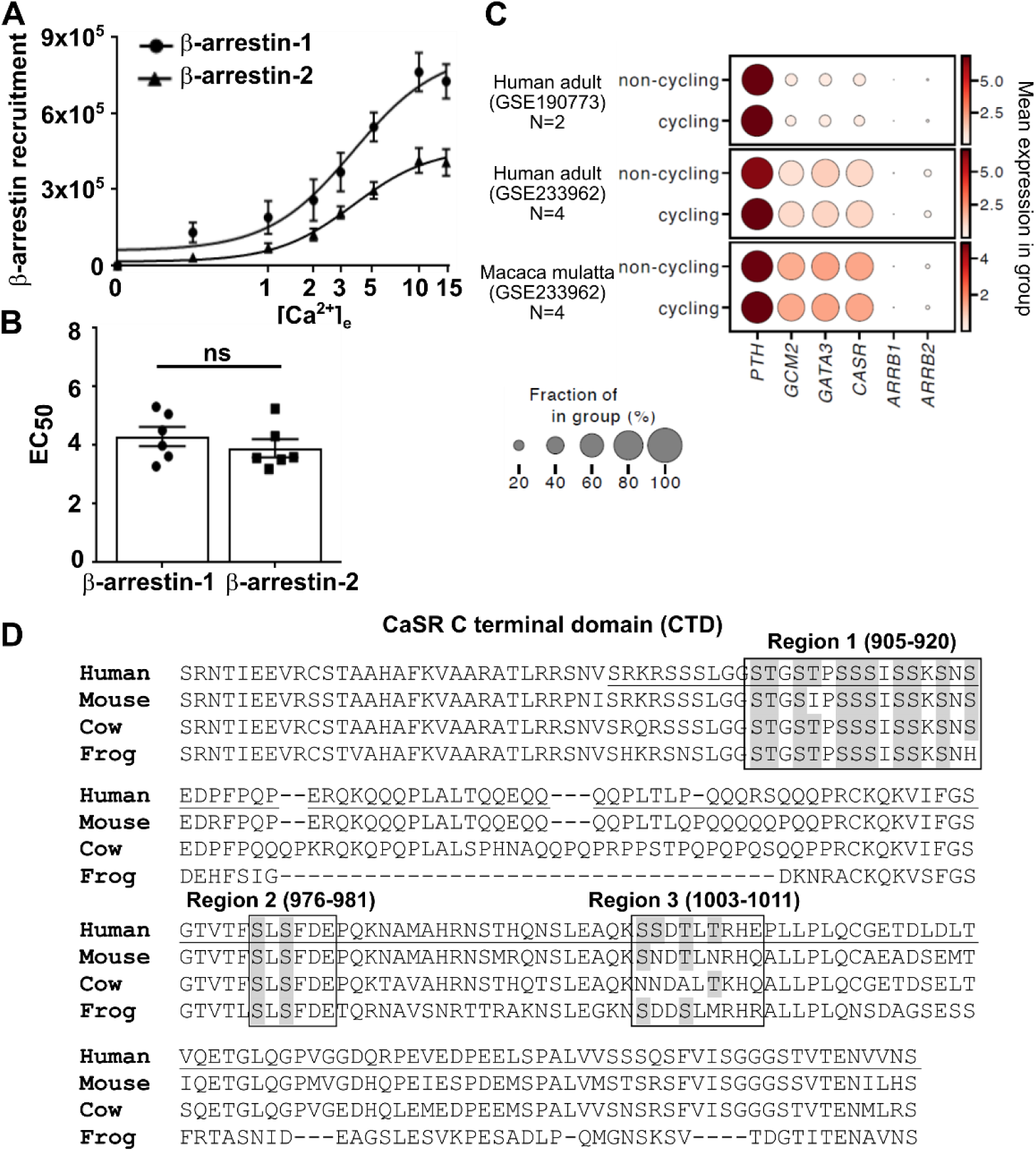
CaSR couples to both β-arrestin-1 and β-arrestin-2 and there are three predicted phosphorylation regions in the C-terminus. (**A**) AUC from luminescence measured in response to increasing concentrations of Ca^2+^_e_ in HEK-βarr1/2 KO cells transfected with CaSR-SmC and either LgC-βarrestin-1 or LgC-βarrestin-2. N=6 biological replicates. Data shows mean±SEM. (**B**) EC_50_ values derived from panel A. (**C**) Dot plot showing the mean expression of canonical parathyroid cell markers (*PTH, GCM2, GATA3, CASR*) together with *ARRB1* and *ARRB2* in parathyroid cells from publicly available datasets (accession numbers indicated next to each panel). Expression levels are shown separately for cycling (G2M and S) and non-cycling (G1) cells. Dot size indicates the proportion of cells within each category with detectable expression. N denotes the number of individuals included from each study. (**D**) Multiple sequence alignment of the CaSR cytoplasmic region with the three predicted phosphorylation sites shown in boxes and the Ser/Thr residues highlighted in gray. Statistical analyses in D were performed by unpaired t-test.

β-arrestin-1 and β-arrestin-2 are expressed in human parathyroids (14) although the relative amounts of each gene have not been assessed. We analyzed three single-cell RNA-sequencing datasets containing non-adenoma parathyroid cells from human adult and rhesus macaque. Expression of β-arrestin-2 was higher in all samples than β-arrestin-1 (Figure 1C) suggesting CaSR is more likely to couple to β-arrestin-2 in parathyroid cells where it performs its primary function.

### The distal cytoplasmic tail of CaSR is required for β-arrestin-1 and β-arrestin-2 recruitment

We used phosphorylation site prediction software to determine possible phosphorylation sites within the cytoplasmic regions of the CaSR that could be important for β-arrestin recruitment. This identified three regions of serine and threonine residues all within the C-terminal tail (Figure 1D). The first region had seven putative phosphorylation sites, although none adhere to the previously described phosphorylation codes used by many GPCRs (3–7,20). The second region between amino acids 976 and 981 (SLSFDE) conforms to a PxPxxP/E/D site (Figure 1D). The third region between amino acids 1003-1011 has four putative phosphorylation sites and similarly conforms to the PxPxxP/E/D site (Figure 1D). We first generated a CaSR plasmid with deletion of part of the cytoplasmic region that encompasses these three putative β-arrestin phosphorylation sites, retains the Thr888 residue that is required for normal CaSR signaling, and replicates mutations identified in individuals with ADH1 in which there is loss of the cytoplasmic region following S895 (28,52) (Figure 2A). The effects of the mutant plasmid were assessed in HEK293 cells depleted of β-arrestin-1/2. The mutant Ser895Stop (895X) CaSR-SmC plasmid was shown to express at cell surfaces similarly to the wild-type plasmid (Figure 2B-C). CaSR-mediated increases in intracellular calcium (Ca^2+^_i_) were significantly greater in cells expressing CaSR-895X when compared to cells expressing the wild-type CaSR plasmid (Figure 2D-E), consistent with previous studies (28,52). The recruitment of β-arrestin-1 and β-arrestin-2 to either CaSR-SmC WT or CaSR-SmC 895X was next assessed in NanoBiT assays. Loss of the distal cytoplasmic region after S895 reduced recruitment of both β-arrestin-1 and β-arrestin-2 (Figure 2F-G) indicating that this region may be important for receptor phosphorylation and subsequent β-arrestin recruitment.

**Figure 2.**
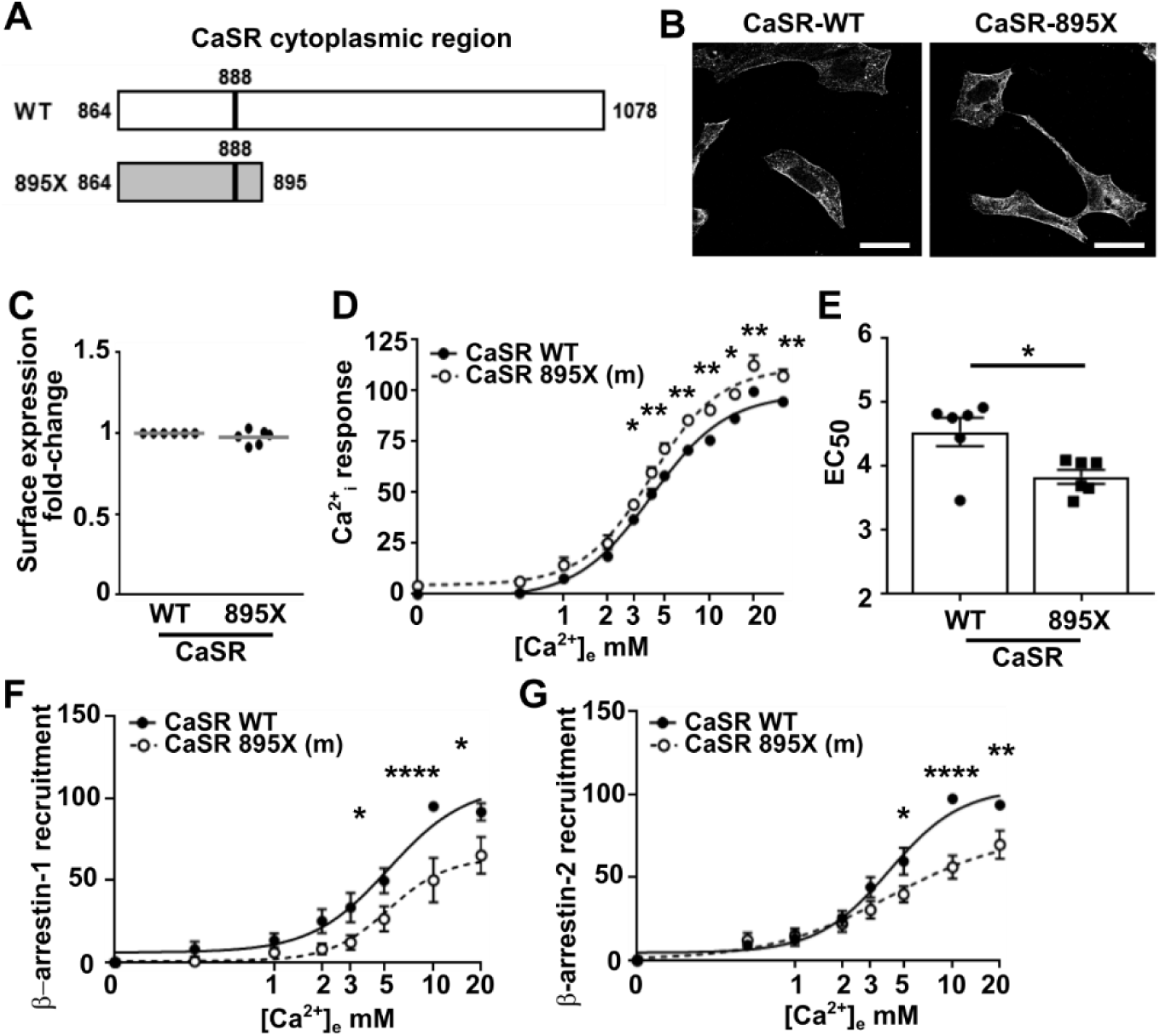
Loss of the distal cytoplasmic region of CaSR enhances signaling and impairs β-arrestin recruitment. (**A**) Cartoon showing the CaSR cytoplasmic region for the WT and 895X truncation plasmid. The Thr888 residue that has previously been shown to have a critical role in CaSR signalling is highlighted and was retained in the truncation plasmid. (**B**) Confocal images of the CaSR-WT and CaSR-895X plasmids in HEK-βarr1/2 KO cells. (**C**) Assessment of cell surface expression measured by labelling surface receptor with a CaSR antibody and a secondary Alexa Fluor 647 antibody. Fluorescence was normalized to cells expressing CaSR-WT. (**D**) Ca^2+^_i_ responses measured by fluo-4 with (**E**) EC_50_ in cells transfected with CaSR WT or 895X. (**F**) βarrestin-1 (N=9) and (**G**) βarrestin-2 (N=9) recruitment to CaSR in cells expressing CaSR WT and 895X plasmids. AUC was measured to derive concentration-response curves. Statistical analyses were performed by two-way ANOVA with Sidak’s multiple-comparisons test. *p<0.05, **p<0.01, ***p<0.001, ****p<0.0001.

### CaSR has distinct phosphorylation sites for β-arrestin-1 and β-arrestin-2 recruitment

We then investigated the Ser and Thr residues within the three regions predicted to have a role in β-arrestin interactions in HEK293 cells depleted of β-arrestin-1/2. The seven serine residues within the first β-arrestin phosphorylation site (S911, S912, S913, S915, S916, S918, S920), two serine residues in the second region (S976, S978) and three residues in the third region (S1004, T1006, T1008) were mutated to alanine, NanoBiT recruitment assays performed and compared to wild-type CaSR responses. Recruitment of β-arrestin-1 was not significantly different to that in cells transfected with wild-type for all residues in the first and second predicted phosphorylation region (Figure 3, Table 1). Mutation of all three residues in the third predicted phosphorylation region reduced β-arrestin-1 recruitment at multiple agonist concentrations. EC_50_ values were significantly greater for the S1004A and T1008A variants (Figure 3, Table 1).

**Figure 3.**
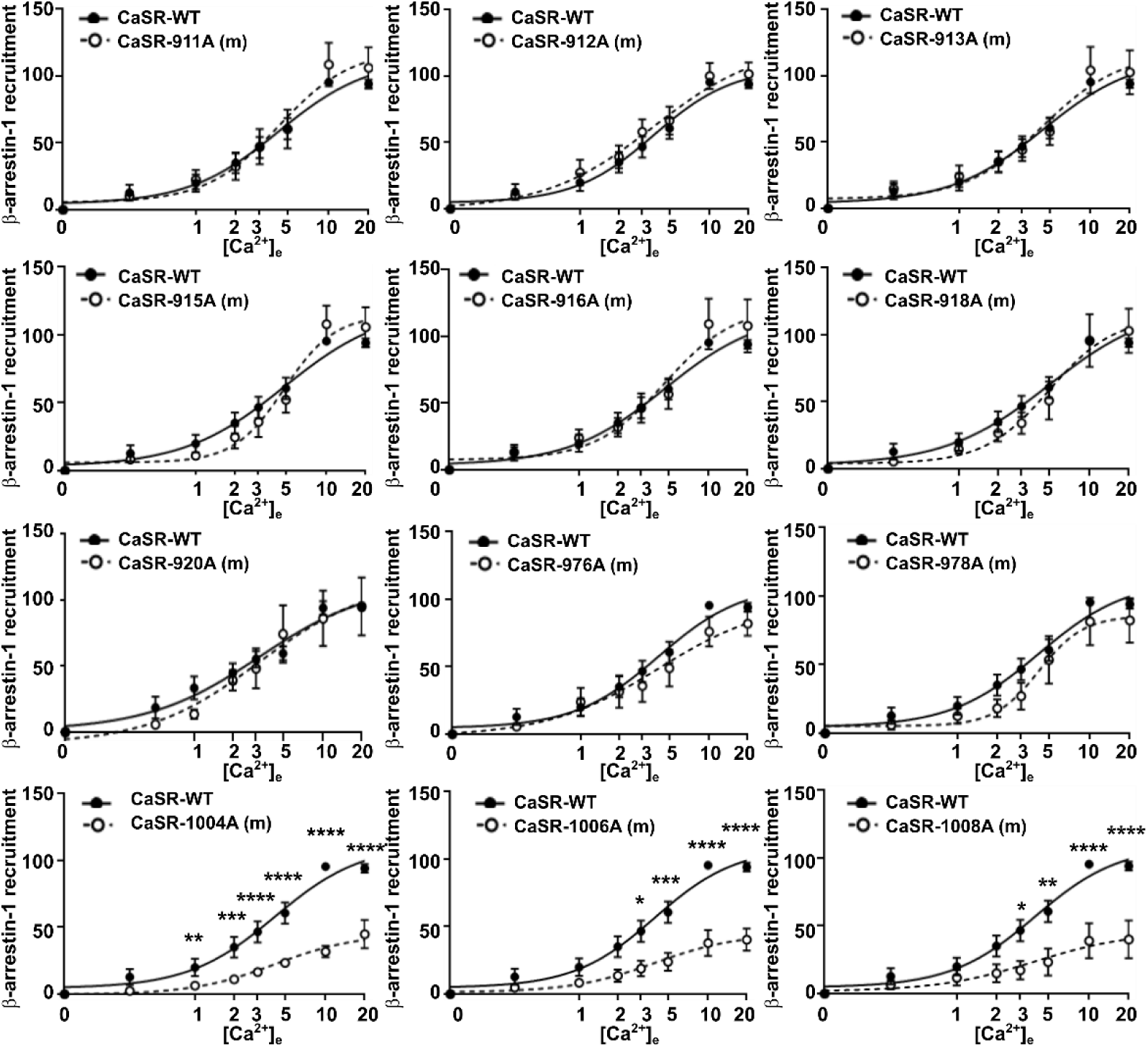
Mutation of residues in the third predicted phosphorylation site impair β-arrestin-1 recruitment to CaSR. β-arrestin-1 recruitment to CaSR in cells expressing CaSR WT and predicted phosphorylation site mutant (m) plasmids. AUC was measured to derive concentration-response curves. Statistical analyses were performed by two-way ANOVA with Sidak’s multiple-comparisons test. N=6-8 biological replicates. *p<0.05, **p<0.01, ***p<0.001, ****p<0.0001.

**Table 1.**
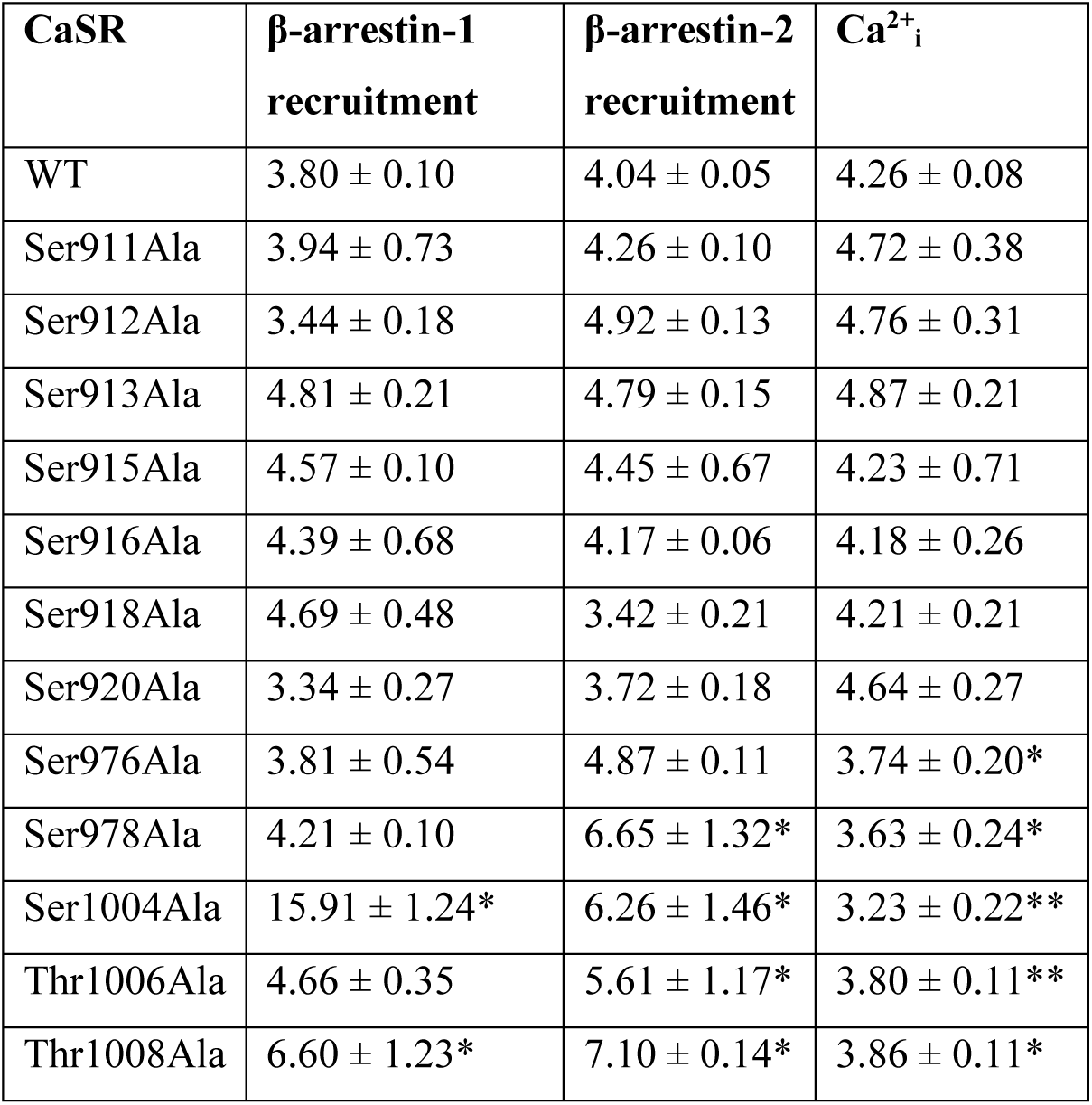
EC_50_ values for β-arrestin-1 and β-arrestin-2 recruitment and Ca^2+^_i_ assays for WT and mutant plasmids. Statistical analyses and pEC_50_ values from concentration-response curves shown in Figure 3-5. Analyses were performed by one-way ANOVA with Dunnett’s multiple-comparisons test and compare each group to WT. *p<0.05, **p<0.01.

NanoBiT recruitment assays were next performed between β-arrestin-2 and the CaSR mutant plasmids. Mutation of six residues (S911, S912, S913, S915, S916, S920) in the first predicted phosphorylation region had little effect on β-arrestin-2 recruitment, although the S918A mutation significantly reduced β-arrestin-2 recruitment without affecting the EC_50_ (Figure 4, Table 1). Mutation of the two residues of the second region reduced both the Emax and enhanced EC_50_ values for β-arrestin-2 recruitment (Figure 4, Table 1). Finally, mutation of the three residues in the third predicted region reduced the Emax and EC_50_ values for β-arrestin-2 recruitment (Figure 4, Table 1). Therefore, mutation of residues in the second and third predicted phosphorylation regions impair both β-arrestin-1 and β-arrestin-2 recruitment, while the second phosphorylation site may also be important for β-arrestin-2 recruitment.

**Figure 4.**
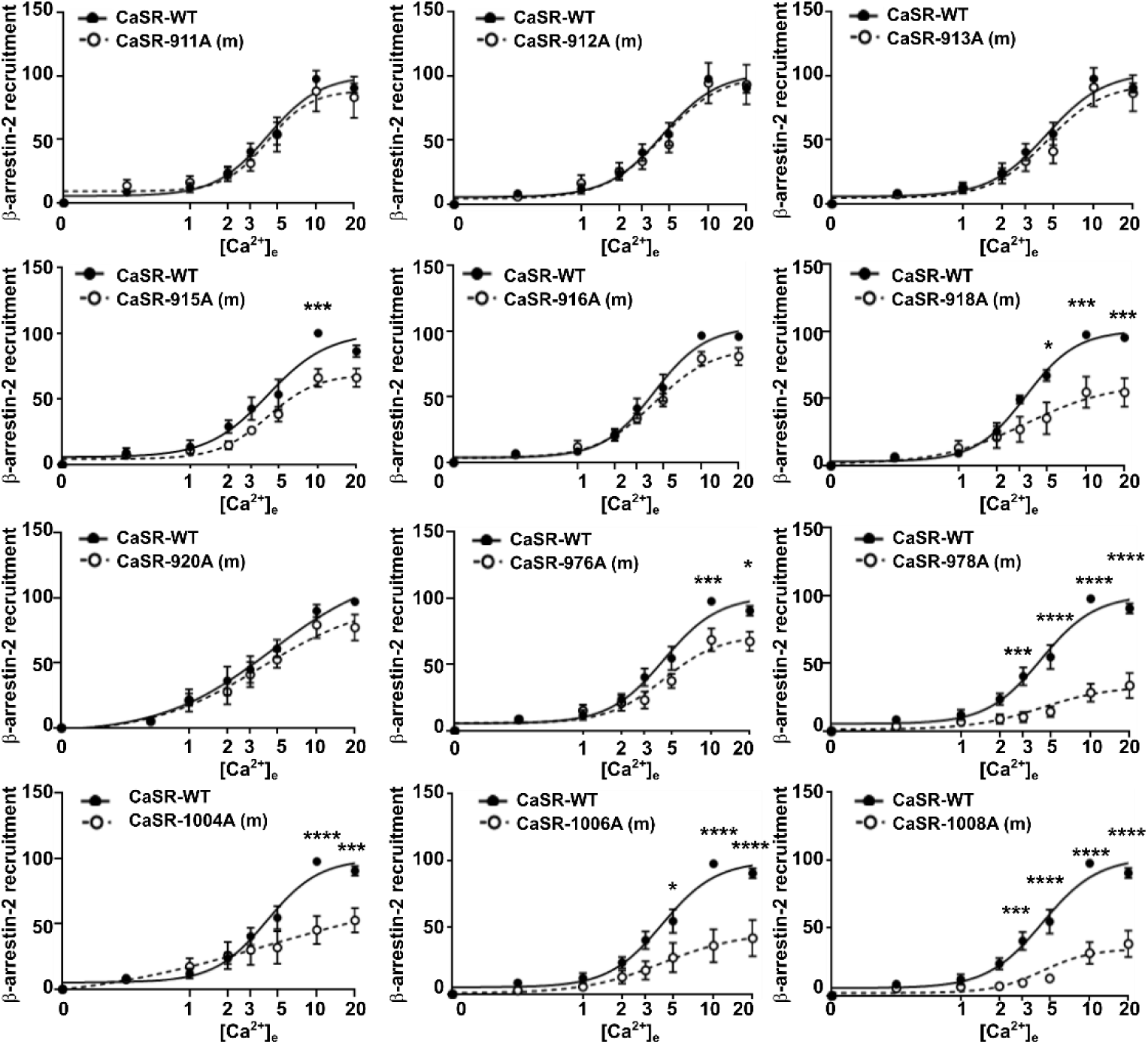
Mutation of residues in the second and third predicted phosphorylation sites impair β-arrestin-2 recruitment to CaSR. β-arrestin-2 recruitment to CaSR in cells expressing CaSR WT and predicted phosphorylation site mutant (m) plasmids. AUC was measured to derive concentration-response curves. Statistical analyses were performed by two-way ANOVA with Sidak’s multiple-comparisons test. N=5-8 biological replicates. *p<0.05, ***p<0.001, ****p<0.0001.

### Mutation of CaSR phosphorylation sites affects receptor signalling

As alanine mutagenesis impaired β-arrestin recruitment, we hypothesized that these mutant proteins would affect CaSR signaling. To assess signaling, CaSR-mediated release of intracellular calcium (Ca^2+^_i_) was measured using fluo-4 assays in cells expressing either CaSR WT or mutant proteins. Signaling for seven residues was not different to cells expressing CaSR WT plasmids (Figure 5A-G). Signaling was enhanced in the two residues of the second predicted phosphorylation site (Figure 5H-I) and the three residues of the third predicted phosphorylation site (Figure 5J-L). Mutation of these five residues significantly reduced EC_50_ values (Table 1), consistent with a gain-of-function. Therefore, mutation of these C-terminal residues impairs β-arrestin recruitment and enhances CaSR downstream signaling.

**Figure 5.**
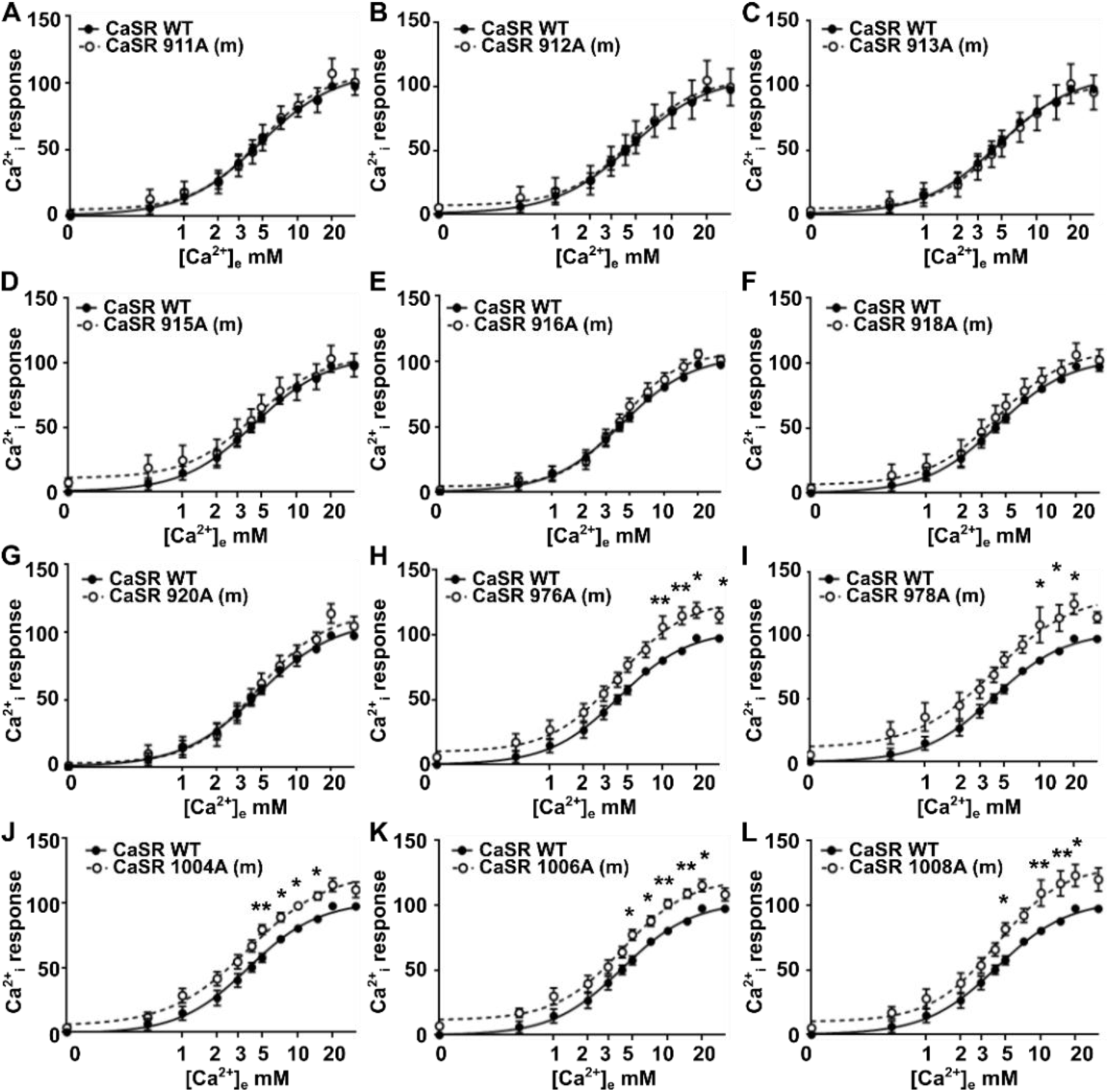
Mutation of residues in the second and third predicted phosphorylation sites of CaSR enhance Ca^2+^_i_ signaling. Normalized Ca^2+^_i_ signaling measured by Fluo-4 assays in cells expressing CaSR WT and predicted phosphorylation site mutant (m) plasmids. The maximal response for each drug concentration was derived and normalized to baseline responses of the CaSR-WT transfected wells to derive the concentration-response curve. Statistical analyses were performed by two-way ANOVA with Sidak’s multiple-comparisons test. N=6-9 biological replicates. *p<0.05, **p<0.01.

### Mutation of the third phosphorylation site affects CaSR internalization

As β-arrestin recruitment was impaired for some residues, we predicted this would reduce CaSR internalization. We generated a CaSR construct with an N-terminal hemagglutinin (HA) epitope followed by a SNAP tag (HA-SNAP-CaSR) to allow tracking of internalization. Expression of the HA-SNAP-CaSR plasmid at the cell surface was confirmed by structured illumination microscopy (SIM) (Figure S1A). The expression of the plasmid was shown to be similar to expression of the previously described FLAG-CaSR construct (49) (Figure S1B). Cells expressing the HA-SNAP-CaSR plasmid had similar CaSR-mediated Ca^2+^_i_ and IP-1 responses to cells expressing the FLAG-CaSR plasmid (Figure S1C-D). Ca^2+^_e_-mediated internalization of the construct was confirmed by SIM imaging (Figure S1E). We then generated alanine mutants of five of the predicted phosphorylation residues in this HA-SNAP-CaSR plasmid, two in site one (915A, 918A), one in site two (978A) and two in site three (1006A, 1008A). Protein expression of the five mutants was similar to the wild-type HA-SNAP-CaSR plasmid (Figure S2A-B), and all mutants were expressed on the cell surface by HILO (Figure S2C). To assess CaSR internalization, cells were exposed to an HA antibody and vehicle or 5mM Ca^2+^_e_ for 30 minutes, then labeled with a fluorescent secondary antibody and the number of internalized vesicles quantified in SIM images. There was a significant difference between the number of vesicles under vehicle conditions (i.e. constitutive internalization) compared to agonist-driven internalization in all groups (Figure 6A-B). However, agonist driven internalization was significantly reduced in cells expressing the 1006A and 1008A mutants compared to cells expressing the CaSR WT plasmid (Figure 6A-B). The basal cell surface expression of CaSR was also significantly greater in cells expressing these two mutant plasmids (Figure 6C).

**Figure 6.**
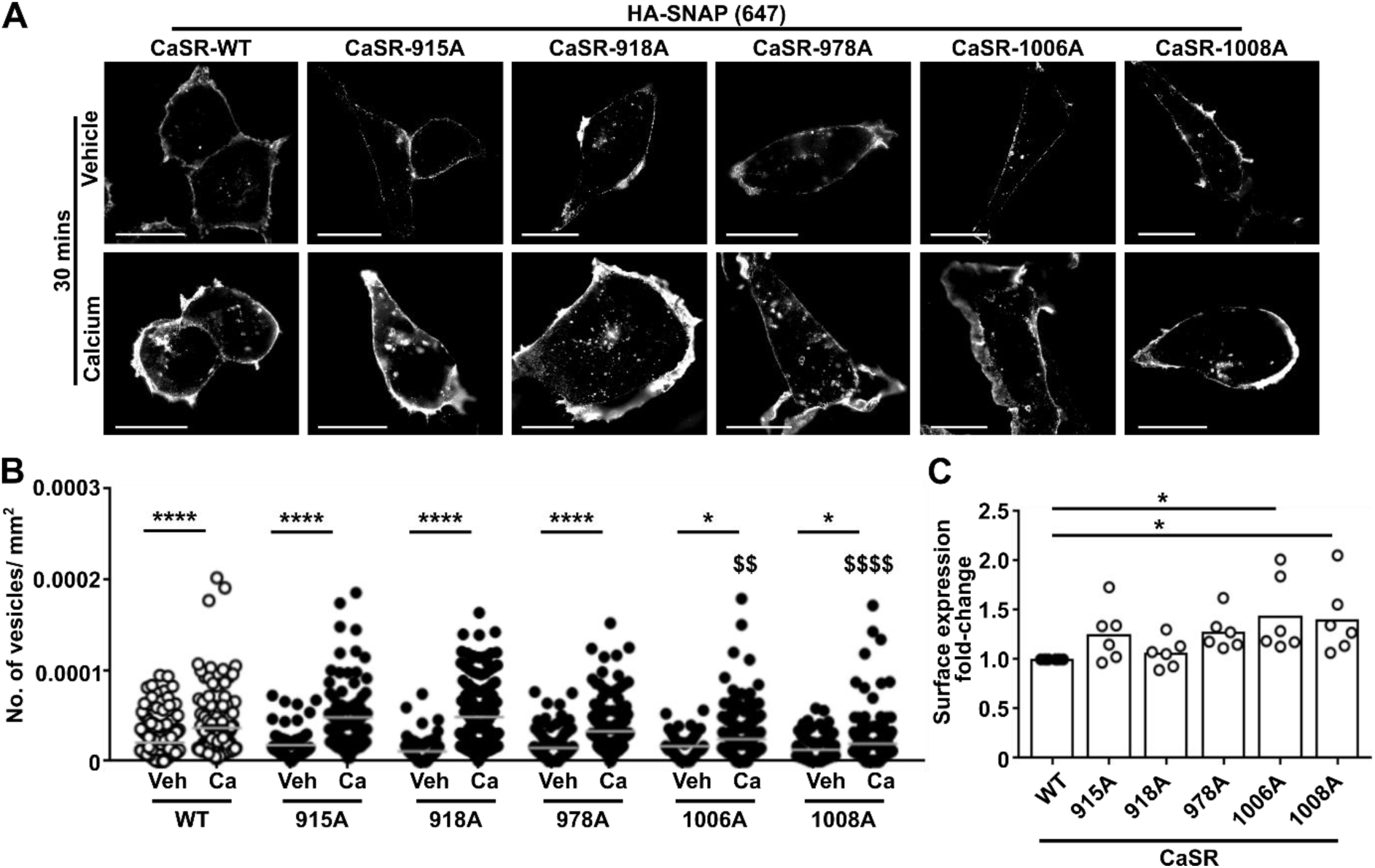
Mutations in the third phosphorylation site impair CaSR internalization. (**A**) SIM of HA-SNAP-CaSR WT and mutant plasmids in AdHEK cells exposed to either vehicle or 5mM Ca^2+^_e_ for 30 minutes. Scale, 5μm. (**B**) Quantification of the number of internalized vesicles. Number of cells from 4-6 independent transfections as follows: WT vehicle (190) and calcium (137), 915A vehicle (70) and calcium (99), 918A vehicle (94) and calcium (159), 978A vehicle (127) and calcium (154), 1006A vehicle (48) and calcium (191), 1008A vehicle (93) and calcium (198). Gray line shows mean. (**C**) Cell surface expression of WT and mutant plasmids. Statistics by one-way ANOVA with Dunnett’s multiple-comparisons test in B and C and compare responses to CaSR-WT. *p<0.05, ***p<0.001, ****p<0.0001. Statistics shown with $ compare agonist responses of each variant with WT calcium responses.

### Rare coding variants impair β-arrestin recruitment and enhance CaSR signaling

Activating mutations in the CaSR are associated with ADH1 and we hypothesized that variants in the CaSR cytoplasmic region identified in individuals undergoing investigation for disorders of calcium homeostasis may be associated with changes in β-arrestin recruitment. No ADH1-associated mutations have been described in the literature, therefore we used the ClinVar database to identify germline genetic variants in *CASR* that have been associated with human disease in the residues that are present in the predicted phosphorylation sites. This identified thirteen variants in total affecting eight different amino acids (Table 2). Most variants in the first phosphorylation site were predicted benign and as alanine mutagenesis of these residues did not affect β-arrestin recruitment, Ca^2+^_i_ signaling or

**Table 2.** CaSR variants located in the cytoplasmic region identified in the ClinVar database. The ClinVar database was accessed on 28/05/24. All variants identified in the predicted phosphorylation sites are shown and annotated with whether they are also present in GnomAD. Pathogenicity was predicted using three tools, CADD (combined annotation dependent depletion)(40), SIFT (sorting intolerant from intolerant)(41) and REVEL (Rare Exome Variant Ensemble Learner). The ClinVar database was accessed on 28/05/24. All variants identified in the predicted phosphorylation sites are shown and whether they are also present in GnomAD listed. Pathogenicity was predicted using three tools, CADD (combined annotation dependent depletion)(40), SIFT (sorting intolerant from intolerant)(41) and REVEL (Rare Exome Variant Ensemble Learner)(42). A CADD score greater than 10 indicates the variant is within the 10% most deleterious substitutions, a score greater than 20 indicates the variant is within the 1% most deleterious substitutions.

| CaSR Variant | Source | Pathogenicity prediction |  |  |
| --- | --- | --- | --- | --- |
|  |  | CADD | SIFT | REVEL |
| **S912F** | ClinVar, GnomAD | 26.3 | predict not tolerated | likely benign |
| **S913F** | ClinVar, GnomAD | 26.3 | predict tolerated | likely benign |
| **S915T** | ClinVar | 26.1 | predict tolerated | likely benign |
| **S920N** | ClinVar, GnomAD | 18.4 | predict tolerated | likely benign |
| **S920R** | ClinVar, GnomAD | 15.2 | predict tolerated | likely benign |
| **S978T** | ClinVar | 26.3 | predict not tolerated | likely benign |
| **S1004R** | ClinVar, GnomAD | 0.481 | predict not tolerated | likely benign |
| **S1004C** | ClinVar | 22.2 | predict not tolerated | likely benign |
| **S1004N** | ClinVar | 0.094 | predict not tolerated | likely benign |
| **T1006A** | ClinVar, GnomAD | 0.407 | predict not tolerated | likely benign |
| **T1006R** | ClinVar | 16.63 | predict not tolerated | likely benign |
| **T1006M** | ClinVar, GnomAD | 17.9 | predict not tolerated | likely benign |
| **T1008P** | ClinVar, GnomAD | 1 | tolerated | likely benign |

CaSR internalization, we did not functionally characterize any ClinVar variants in these residues. There were no variants in S978 and only a single variant in S978 to a T978 (Table 2). As this variant residue can still be phosphorylated, we chose not to investigate it any further. There were seven variants in the third predicted phosphorylation site (S1004R/C/N, T1006A/R/M, T1008P) and we chose one variant from each residue to characterize in further detail. The CaSR-SmC plasmid was mutated to either 1004C, 1006M or 1008P and β-arrestin recruitment assays performed. The CaSR 1004C variant had no effect on β-arrestin-1 or β-arrestin-2 recruitment (Figure 7A-B, Table 3). In contrast, the 1006M and 1008P variants reduced β-arrestin-1 and β-arrestin-2 recruitment to CaSR, with reduced EC_50_ for 1006M (Figure 7A-B, Table 3). Ca^2+^_i_ recruitment was significantly elevated in cells expressing the 1006M and 1008P variants, with reduced EC_50_ for 1006M, consistent with a gain-of-functional activity (Figure 7C, Table 3). Ca^2+^_i_ signaling in cells expressing the 1004C variant was similar to cells expressing CaSR WT.

**Figure 7.**
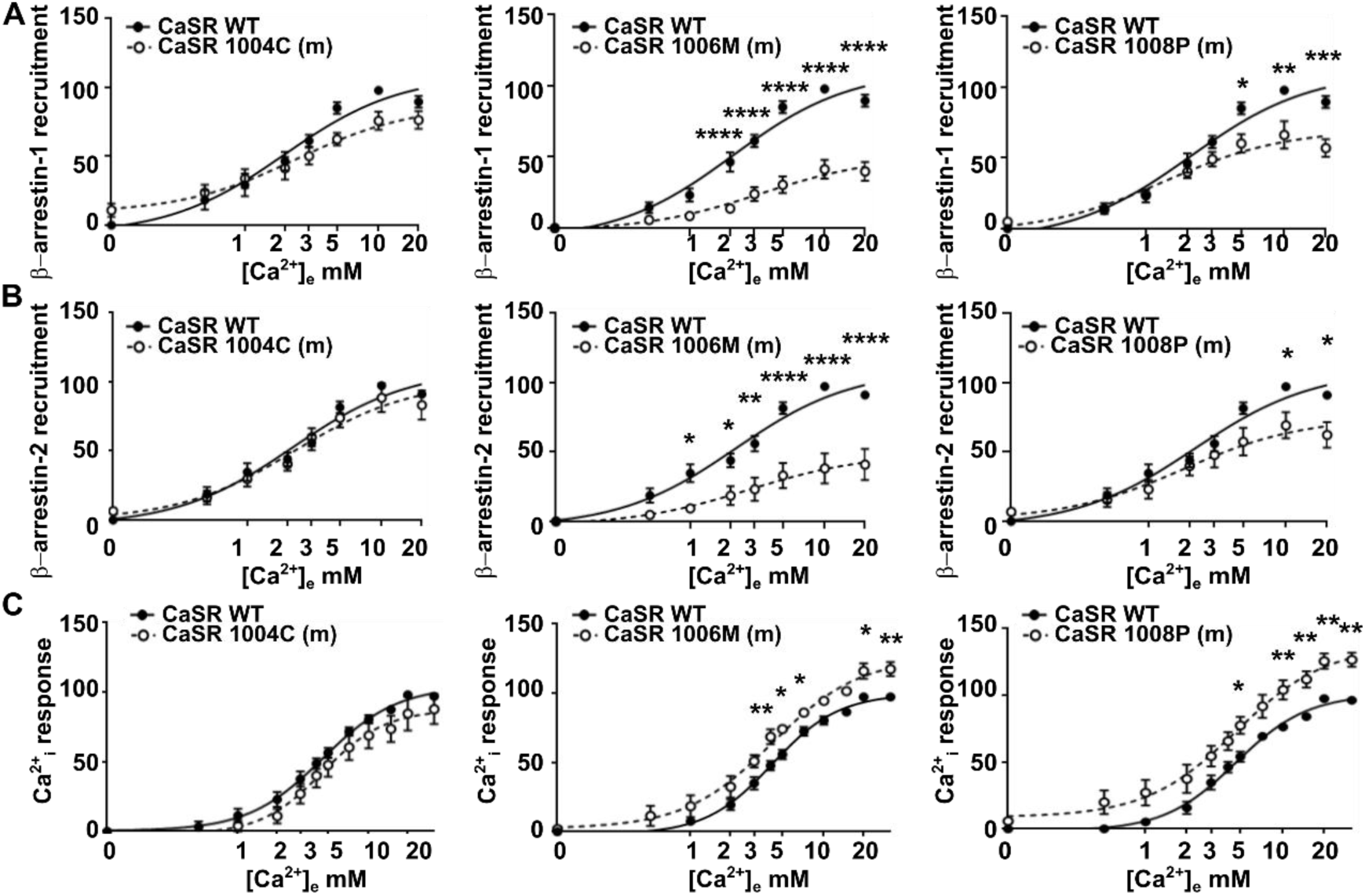
ClinVar mutants in the third phosphorylation site impair β-arrestin recruitment and enhance CaSR signaling. (**A**) β-arrestin-1 recruitment (N=5) and (**B**) β-arrestin-2 recruitment (N=7) to CaSR WT and plasmids with ClinVar mutants (m) in three residues of the third phosphorylation site. AUC was measured to derive concentration-response curves. (**C**) Normalized Ca^2+^_i_ signaling measured by Fluo-4 assays in cells expressing CaSR WT and plasmids with ClinVar mutants (m). N=6-7 biological replicates. The maximal response for each drug concentration was derived and normalized to baseline responses of the CaSR-WT transfected wells to derive the concentration-response curve. Statistical analyses were performed by two-way ANOVA with Sidak’s multiple-comparisons test.

**Table 3.** EC_50_ values for β-arrestin-1 and β-arrestin-2 recruitment and Ca^2+^_i_ assays for ClinVar mutants. Statistical analyses and pEC_50_ values from concentration-response curves shown in Figure 7. Analyses were performed by one-way ANOVA with Dunnett’s multiple-comparisons test and compare each group to WT. *p<0.05, **p<0.01.

| CaSR | β-arrestin-1<br>recruitment | β-arrestin-2<br>recruitment | $Ca^{2+}_i$ |
| --- | --- | --- | --- |
| WT | $2.14 \pm 0.31$ | $2.14 \pm 0.40$ | $4.33 \pm 0.26$ |
| Ser1004Cys | $2.06 \pm 0.39$ | $2.29 \pm 0.37$ | $4.58 \pm 1.07$ |
| Thr1006Met | $3.38 \pm 0.38^*$ | $3.33 \pm 0.14^*$ | $3.59 \pm 0.12^*$ |
| Thr1008Pro | $1.49 \pm 0.18$ | $1.87 \pm 0.38$ | $4.33 \pm 0.67$ |

To determine whether these changes in signaling also affected internalization as we observed for the phosphorylation site mutants (Figure 6), we generated a HA-SNAP-CaSR-1006M plasmid and performed SIM imaging. The number of internalized vesicles in cells expressing the 1006M plasmid was significantly reduced compared to WT expressing cells following exposure to vehicle or calcium for 30 minutes (Figure 8A-B), indicating that both agonist-driven and constitutive internalization was impaired in cells with the ClinVar mutant. There was no significant difference in cell surface expression for the CaSR WT or 1006M variant under basal conditions (Figure 8C). As MAPK signaling has been reported downstream of β-arrestin, we also examined the phosphorylation of ERK1/2 (pERK1/2) in response to the 1006M ClinVar and the 1006A phosphorylation site mutants. Cells were exposed to Ca^2+^_e_ for 5 and 10 minutes and pERK1/2 levels assessed by western blot analysis. When pERK1/2 concentrations were normalized to total ERK1/2 levels, there was no significant difference detected for either the 1006M or 1006A mutants (Figure 8D-G, S3A-B). Therefore, CaSR cytoplasmic region variants in the ClinVar database impair β-arrestin recruitment and enhance CaSR Ca^2+^_i_, but do not affect pERK1/2 signaling.

**Figure 8.**
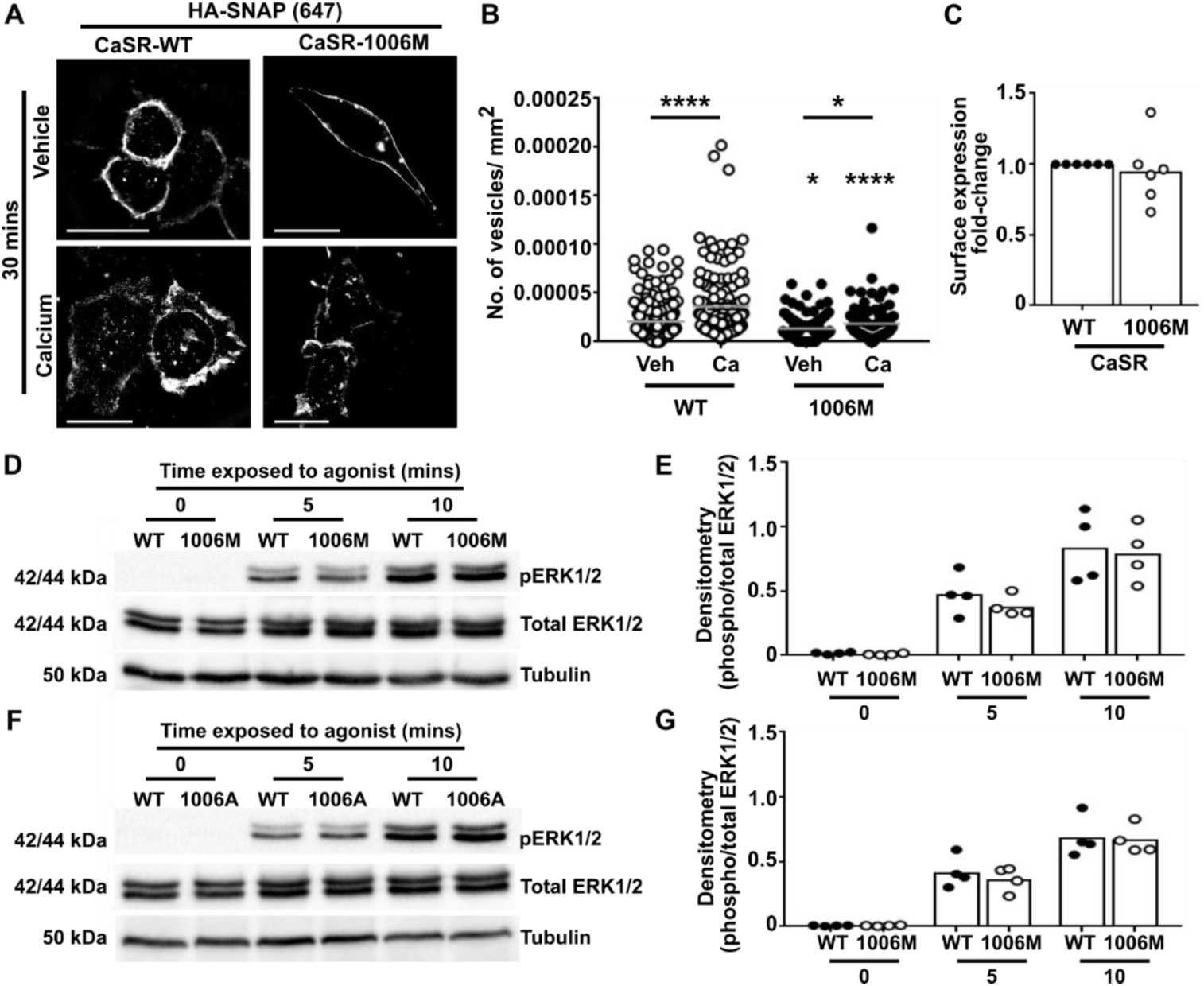
The CaSR-1006M ClinVar mutant impairs internalization but has normal pERK1/2 signaling. (**A**) SIM of HA-SNAP-CaSR WT and the 1006M ClinVar mutant plasmid in AdHEK cells exposed to either vehicle or 5mM Ca^2+^_e_ for 30 minutes. Scale, 5μm. (**B**) Quantification of the number of internalized vesicles. Number of cells from 6-7 independent transfections as follows: WT vehicle (190) and calcium (137), 1006M vehicle (119) and calcium (92). Gray line shows mean. (**C**) Assessment of cell surface expression of CaSR WT and the 1006M mutant. Statistics by one-way ANOVA with Dunnett’s multiple-comparisons test and compare responses to CaSR-WT in B and C. (**D**) Western blot analysis of phosphorylated ERK1/2 (pERK1/2) in cells expressing HA-SNAP-CaSR WT and 1006M ClinVar mutant in response to 5mM Ca^2+^_e_. (**E**) Densitometry analysis of pERK1/2 normalized to total ERK1/2 in four blots shown in Figure S3. (**F**) Western blot analysis of phosphorylated ERK1/2 (pERK1/2) in cells expressing HA-SNAP-CaSR WT and 1006A mutant in response to 5mM Ca^2+^_e_. (**G**) Densitometry analysis of pERK1/2 normalized to total ERK1/2 in four blots shown in Figure S3 *p<0.05, **p<0.01, ****p<0.0001.

## Discussion

Our studies demonstrated that the CaSR CTD contains multiple residues that are important for β-arrestin recruitment. Regions in the CTD were predicted, using computational tools, to conform to partial or full phosphorylation sites, and a truncation mutation that resulted in the loss of all three predicted sites impaired β-arrestin recruitment and enhanced receptor signaling. We used alanine mutagenesis of individual residues within these sites to show that β-arrestin recruitment was impaired by introduction of non-phosphorylated residues within two of these sites. These latter two sites conform to previously described phosphorylation codes of the PxPxxP/E/D format (6). While alanine mutagenesis of three residues in the third phosphorylation site impaired both β-arrestin-1 and β-arrestin-2 recruitment, mutation of S976 and S978 of site 2 only impaired β-arrestin-2 recruitment. This may be because each arrestin protein recognises different phosphorylation codes and therefore may bind in different conformations to the receptor, as was described for the parathyroid hormone type-1 receptor (PTH1R)(53) and the vasopressin receptor (V_2_R)(54). Alternatively, these different codes may be recognised by distinct GRKs, which could bind different forms of β-arrestin, similar to that described to encode differential functions of β-arrestin downstream of the β2-adrenergic receptor (3). Previous studies have shown that GRK4 overexpression inhibits CaSR signaling by enhancing phosphorylation and β-arrestin translocation, while GRK2 overexpression disrupts G_q_ signaling by phosphorylation-independent mechanisms (14). Thus, there is precedent for differential effects mediated by GRKs at the CaSR, and future studies could focus on how different GRK proteins are recruited to each of the CaSR phosphorylation sites. Irrespective of this difference in β-arrestin recruitment, CaSR response to Ca^2+^_i_ was enhanced.

We showed that CaSR is capable of recruiting β-arrestin-1 and β-arrestin-2, consistent with previous findings that the receptor interacts with both proteins (14), and that β-arrestin recruitment reduces signaling consistent with the classical model of GPCR desensitization (14–16). Our analysis of single-cell RNA sequencing datasets showed that β-arrestin-2 expression is higher than β-arrestin-1 and it is therefore likely that CaSR predominantly recruits β-arrestin-2 in physiologically relevant cells. One previous study was unable to identify a role for β-arrestin in CaSR signaling (17); however, they focussed on the ERK1/2 pathway which we similarly showed was not affected by mutation of phosphorylation sites that did impair β-arrestin recruitment. Additionally, signaling of CaSR from intracellular sites has also been shown to be β-arrestin-independent (12). Collectively, these studies suggest CaSR can perform some signaling in the absence of β-arrestin, similarly to other class C GPCRs, including the GABA_B_ receptor (55) and some mGluRs (23).

We also showed that mutation of the third predicted phosphorylation motif, but not residues in the other motifs, affected CaSR agonist-driven internalization. Mutations within this motif affect both β-arrestin-1 and β-arrestin-2 binding, suggesting this motif may have a more important role in receptor internalization than the second motif. Studies of the V_2_R have shown some phosphorylation sites are more important for trafficking than others (54). Although some previous studies of the CaSR suggested that β-arrestin recruitment had no role in CaSR internalization (14,15), a more recent study that used real-time assays showed β-arrestins did contribute to CaSR internalization (19). It is possible that the earlier studies missed this effect as they used cell surface expression as a readout for internalization (14,15). Alternatively, it may be because CaSR has considerable constitutive internalization (2,12,19,56) that could have masked agonist-driven effects in these previous studies. Constitutive internalization was not affected by mutation of any of the residues. This contrasts with a previous study that characterized CaSR internalization in HEK293 cells depleted of β-arrestin-1 and β-arrestin-2, which showed both reduced constitutive and agonist-driven internalization (19). However, our assays only looked at the effect of mutation of single residues, not mutation of the whole predicted phosphorylation site, and it is likely that there would be an additive effect of the mutations that would be more analogous to complete depletion of β-arrestin proteins. Previous studies of phosphorylation barcodes have shown such additive effects on the affinity of interactions (5,7,54).

Some additional differences were detected between β-arrestin-1 and β-arrestin-2 recruitment to the CaSR. Luciferase values for β-arrestin-1 were higher than β-arrestin-2, which could suggest β-arrestin-1 forms tighter associations with the receptor. However, there was no difference in the EC_50_ values between the two recruitment assays, and it is possible that these differences are due to the position of the LgBiT tag on the β-arrestin proteins, which may not be optimal for coupling to CaSR. Other measures of protein affinity would need to be performed to definitively characterize this. This is particularly important, as all studies that have assessed receptor affinity have demonstrated β-arrestin-2 has a higher affinity than β-arrestin-1 (57).

Deletion of the distal CTD of the CaSR impaired β-arrestin recruitment and impaired receptor signaling. Previous studies that also examined truncation of the CaSR CTD encompassing this region also identified enhanced signaling, attributed to loss of degradation motifs (24,34). Our studies suggest the enhanced signaling observed in these studies could also be due to impaired β-arrestin recruitment. Thus, ADH mutations due to truncations within this region (28,29) could be associated with receptor activation due to loss of degradation motifs (24,34), impaired ability to recruit β-arrestin, and loss of an internalization motif. Based on these studies, we hypothesized that other clinically relevant variants in the CaSR may affect β-arrestin recruitment and internalization and that this could modify receptor signaling. However, most disease-associated mutations in the CaSR affect residues within the extracellular domain or transmembrane regions (1), and we instead selected variants from the ClinVar database that have been associated with disorders of calcium homeostasis. A caveat of using ClinVar is that both activating and inactivating mutations in the CaSR are reported as ‘disorders of calcium homeostasis’ and it is not possible to determine whether individuals with these variants are hypo- or hypercalcemic. We assessed three variants affecting residues in the third predicted phosphorylation site and showed that two impaired β-arrestin recruitment and enhanced receptor signaling. Further characterization of one of these variants demonstrated impaired internalization, consistent with the reduction in β-arrestin recruitment and enhanced CaSR signaling. This suggests that some CaSR variants may affect receptor signaling by modifying receptor phosphorylation and β-arrestin recruitment, similar to disease-causing variants in other GPCRs (58,59). As CaSR is an obligate dimer it is possible that it may engage multiple β-arrestin proteins simultaneously as was shown for mGluRs (60), and therefore the impact of heterozygote mutations is unknown. Investigation of the calcium levels in individuals with these variants would be useful to determine the relative effects of impaired β-arrestin recruitment, although a large population study may be required to obtain sufficient subjects to determine associations.

In conclusion, our studies have identified two regions within the CaSR CTD that are important for recruitment of β-arrestin and affect downstream signaling pathways. Clinically-relevant mutations that abolish or disrupt these sites are associated with impaired β-arrestin recruitment and enhanced CaSR signaling and this may represent a novel mechanism by which some ADH mutations cause hypocalcemia.

## Declaration of Interest

The authors have no conflicts of interest to declare.

## Funding

This work was supported by: An Academy of Medical Sciences Springboard Award supported by the British Heart Foundation, Diabetes UK, the Global Challenges Research Fund, the Government Department of Business, Energy and Industrial Strategy and the Wellcome Trust (Ref: SBF004|1034, C.M.G) and a Sir Henry Dale Fellowship jointly funded by the Wellcome Trust and the Royal Society (Grant Number 224155/Z/21/Z to CMG). HM was supported by a Blavatnik Fellowship (British Council) and The Israeli Council for Higher Education.

## Author contributions

Conceptualization: CMG

Methodology: HM, CMG

Investigation: RAW, HM, CMG

Writing – original draft: CMG

Writing – review and editing: RAW, HM, CMG

## Data and materials availability

All data needed to evaluate the conclusions in the paper are present in the paper and/or the Supplementary Materials. Raw data will be made available upon request. Plasmid constructs developed for this manuscript will readily be made available upon request. Plasmid constructs obtained from other researchers are detailed in the methods section and may be subject to Material Transfer Agreements. Please contact the corresponding author of this manuscript, or the named source for details.

## Supplementary Appendix

**Figure S1.**
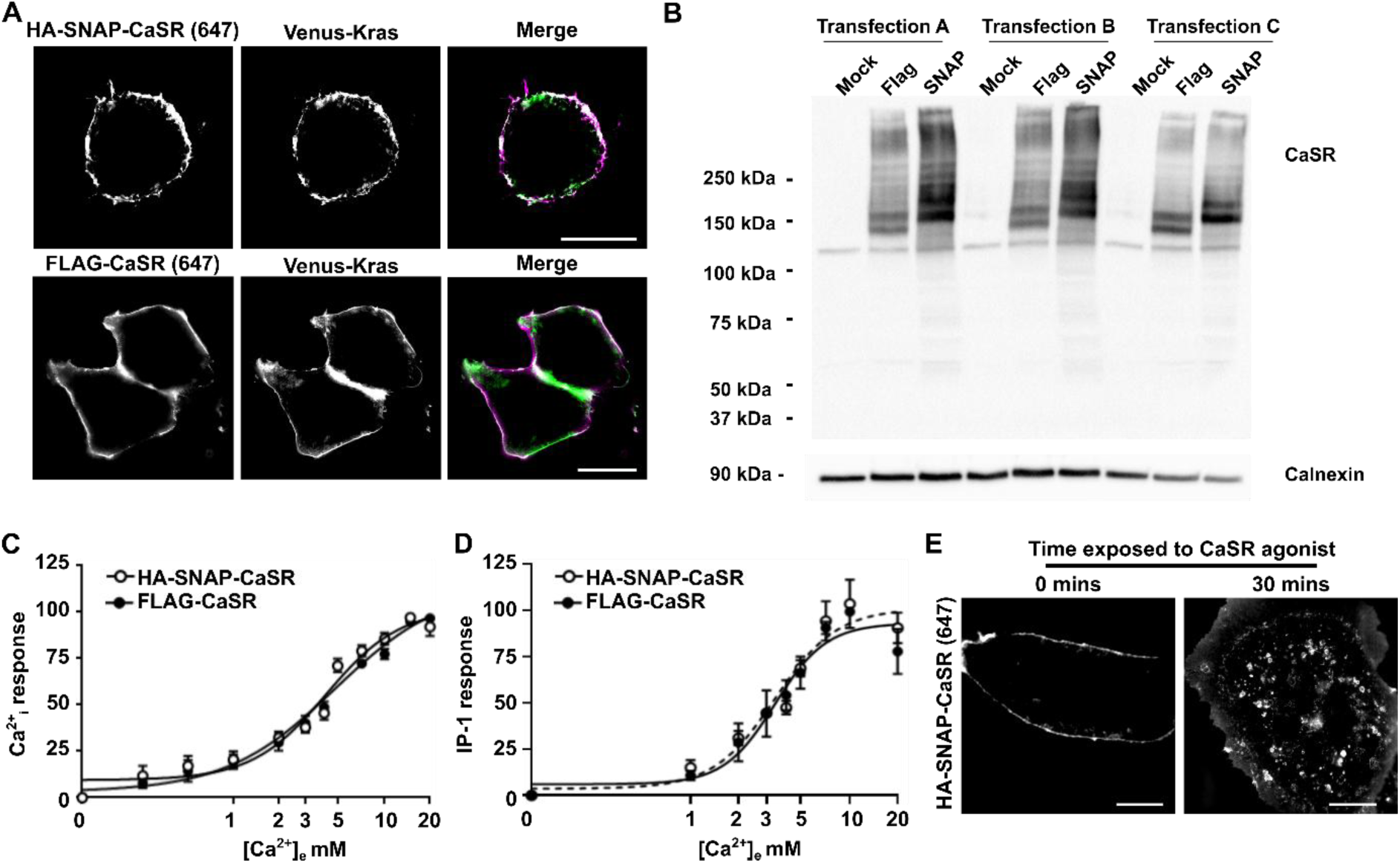
Generation of the HA-SNAP-CaSR plasmid. (**A**) SIM of HA-SNAP-CaSR (labelled with the HA antibody and an Alexa Fluor 647 secondary antibody) and Venus-Kras in AdHEK cells. Scale, 5μm. (**B**) Western blot analyses to detect CaSR in cells transfected with mock (pcDNA3.1), FLAG-CaSR-WT (FLAG) and HA-SNAP-CaSR-WT (CaSR) plasmids showing similar expression levels. Calnexin was used as a loading control. The blot shows lysates from three independent transfections. (**C**) Normalized Ca^2+^_i_ signaling measured by Fluo-4 assays. N=5 biological replicates. The maximal response for each drug concentration was derived and normalized to baseline responses of the CaSR-WT transfected wells to derive the concentration-response curve. (**D**) IP-1 responses in cells transfected with HA-SNAP-CaSR and FLAG-CaSR. N=3 biological replicates. (**E**) SIM images of HA-SNAP-CaSR exposed to 5mM Ca^2+^_e_ and the HA antibody for 0 or 30 minutes.

**Figure S2.**
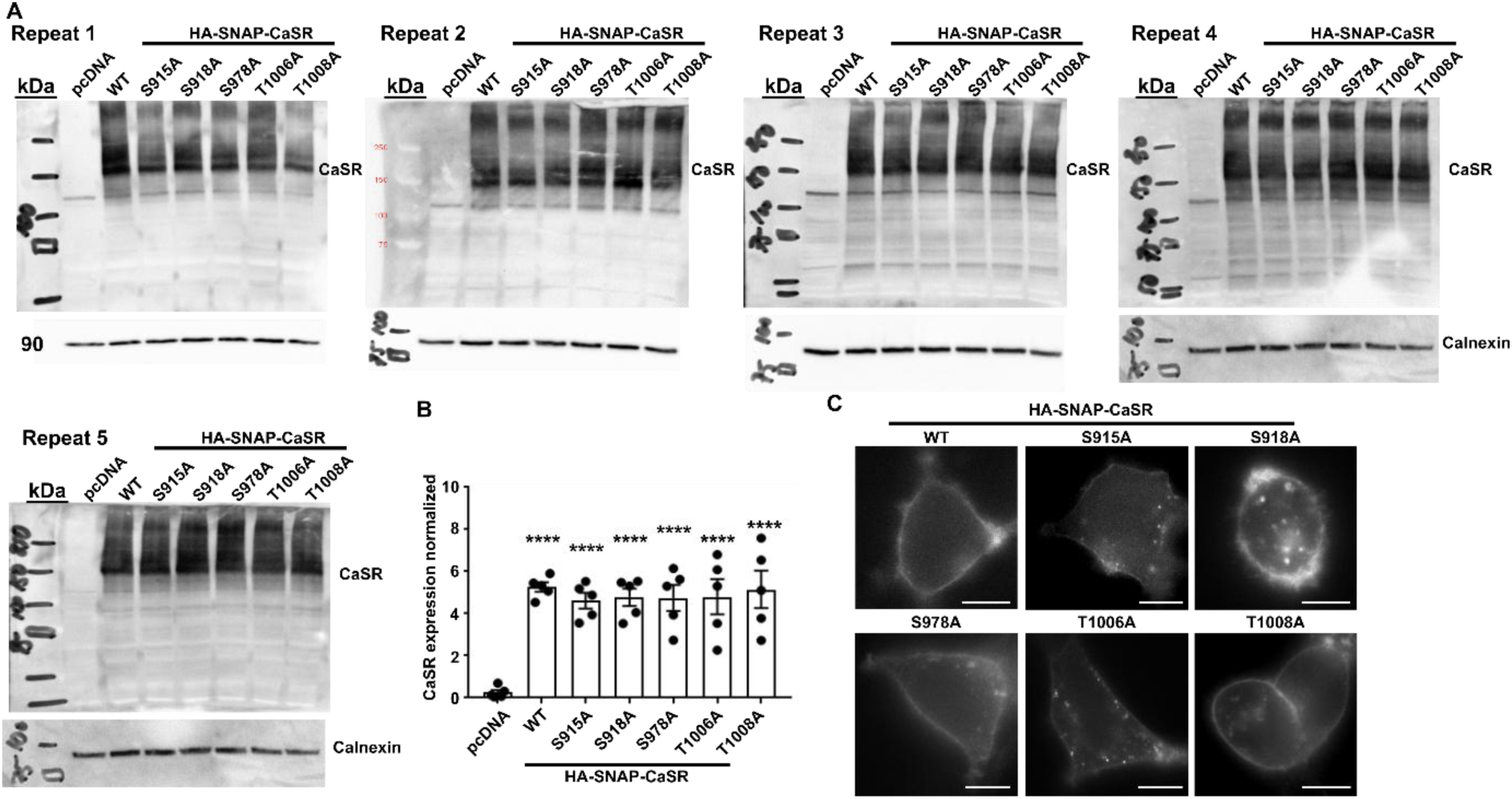
Total protein expression and surface expression of the HA-SNAP-CaSR WT and mutant plasmids. (**A**) Western blot analyses to detect CaSR in cells transfected with mock (pcDNA3.1), HA-SNAP-CaSR-WT and mutant plasmids. (**B**) Densitometry of CaSR expression from the five blots normalized to calnexin as a loading control. N=5 biological replicates. Statistical analyses were performed by one-way ANOVA with Dunnett’s multiple-comparisons test. (**C**) HILO imaging of HA-SNAP-CaSR WT and mutant plasmids showing the receptor is expressed at the cell surface. Scale, 5μm.

**Figure S3.**
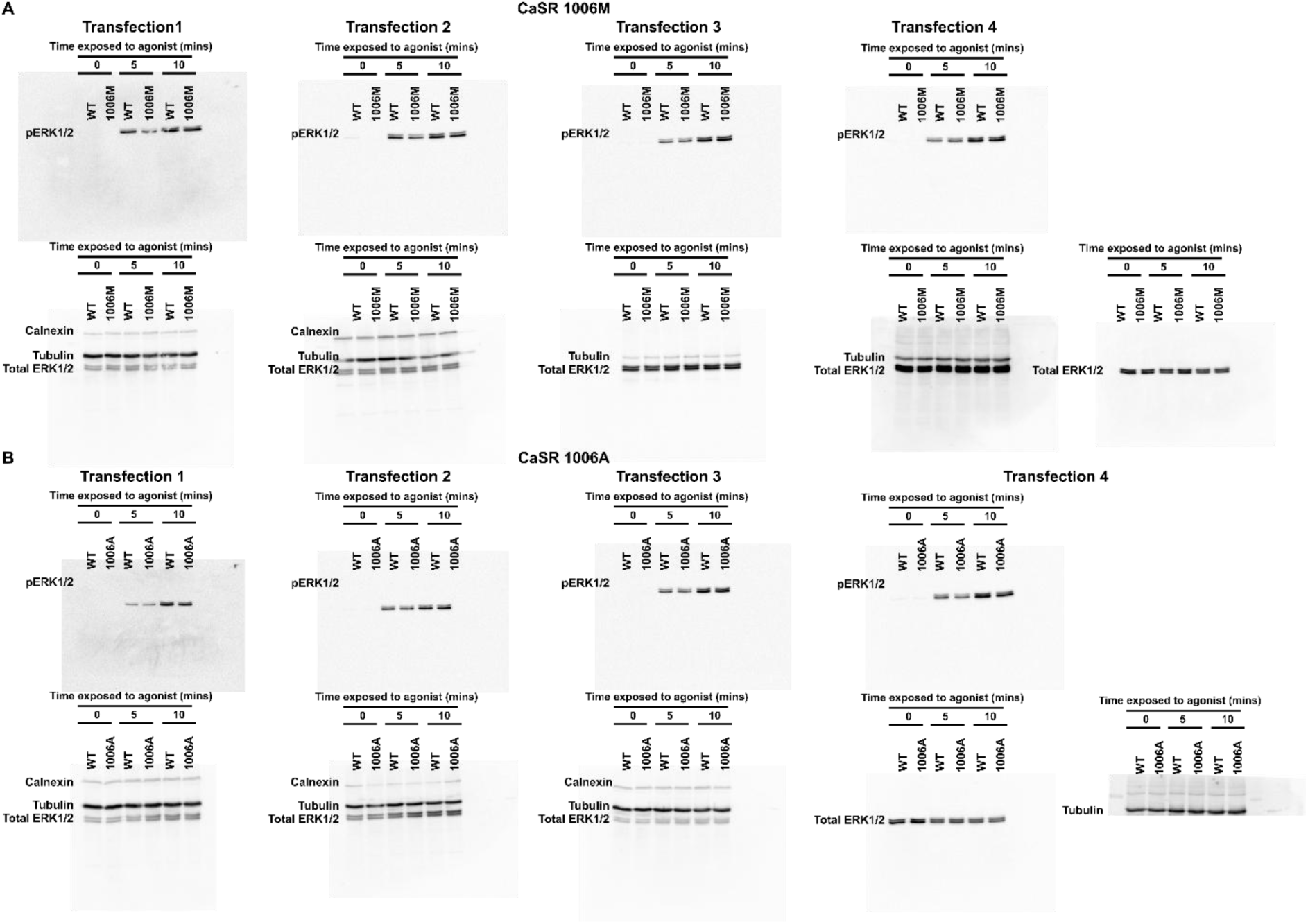
Whole blots for pERK1/2 densitometry shown in Figure 8. (**A**) Full blots for westerns shown in Figure 8D and (**B**) Figure 8F.

## Notes

### Competing Interest Statement

The authors have declared no competing interest.

